# Mutation, selection, and the prevalence of the *C. elegans* heat-sensitive mortal germline phenotype

**DOI:** 10.1101/2021.11.29.470498

**Authors:** Sayran Saber, Michael Snyder, Moein Rajaei, Charles F. Baer

## Abstract

*C. elegans* strains with the heat-sensitive mortal germline (Mrt) phenotype become progressively sterile over the course of a few tens of generations when maintained at temperatures near the upper range of *C. elegans*’ tolerance. Mrt is transgenerationally-heritable, and proximately under epigenetic control. Previous studies have suggested that Mrt presents a relatively large mutational target, and that Mrt is not uncommon in natural populations of *C. elegans*. The Mrt phenotype is not monolithic. Some strains exhibit a strong Mrt phenotype, in which individuals invariably become sterile over a few generations, whereas other strains show a weaker (less penetrant) phenotype in which the onset of sterility is slower and more stochastic. We present results in which we (1) quantify the rate of mutation to the Mrt phenotype, and (2) quantify the frequency of Mrt in a collection of 95 wild isolates. Over the course of ~16,000 meioses, we detected one mutation to a strong Mrt phenotype, resulting in a point estimate of the mutation rate *U*_*Mrt*_**≈** 6×10^−5^/genome/generation. We detected no mutations to a weak Mrt phenotype. 6/95 wild isolates have a strong Mrt phenotype, and although quantification of the weak Mrt phenotype is inexact, the weak Mrt phenotype is not rare in nature. We estimate a strength of selection against mutations conferring the strong Mrt phenotype 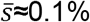, similar to selection against mutations affecting competitive fitness. The appreciable frequency of weak Mrt variants in nature combined with the low mutation rate suggests that Mrt may be maintained by balancing selection.

## Introduction

The *C. elegans* “mortal germline” (Mrt) phenotype is a transgenerationally heritable trait in which Mrt lineages become progressively sterile over the course of a few to a few tens of generations (Ahmed and Hodgkin 2000; Smelick and Ahmed 2005). The Mrt phenotype was first discovered in a mutant strain defective in germline telomere replication and DNA repair (Ahmed and Hodgkin 2000). Subsequent studies have identified numerous Mrt mutants, many of which are associated with defects in nuclear RNAi (Katz et al. 2009; Buckley et al. 2012; Spracklin et al. 2017). The transgenerational heritability of the nRNAi-defective Mrt phenotype is under proximate epigenetic control, often (perhaps always) involving the interplay between piRNAs and their Argonaute protein partner *prg-1* (Batista et al. 2008; Wahba et al. 2021). However, like any trait governed epigenetically, it has an ultimate, underlying genetic basis. nRNAi-defective Mrt mutants are typically heat-sensitive, with continued exposure to high temperature leading to onset of sterility. Based on the frequency of appearance of Mrt mutants in forward genetic screens, it was suggested that many genes are capable of producing the Mrt phenotype if mutated (Smelick and Ahmed 2005), or in other words, the Mrt phenotype presents a large mutational target (Houle 1998). Note that "large mutational target" in this context is not synonymous with "polygenic" in the usual sense, because even if many genes potentially affect the trait, a mutation in any one gene is sufficient to produce the Mrt phenotype. On the other hand, Mrt probably is polygenic, with subtle phenotypic variation resulting from segregating variants at many loci. However, it would be challenging to discern whether a given genotype becomes sterile after (say) 13 generations vs. 14 generations, on average.

Over the past two decades, the realm of *C. elegans* biology has expanded beyond its initial role as a model system par excellence for functional biology to include studies of natural variation (Dirksen et al. 2016; Felix and Duveau 2012; Schulenburg and Felix 2017; Cook et al. 2017). It soon became apparent that some wild isolates could not be maintained in culture at 25° C (near the upper range of *C. elegans* thermal tolerance), and further, that most such strains had the heat-sensitive Mrt phenotype (Frezal et al. 2018). Heat-sensitive Mrt strains can typically be rescued by exposure to cool temperature (15° C) for a generation, and it is unclear if long-term (multi-generational) exposure to temperatures sufficiently high to induce the Mrt phenotype is common in *C. elegans’* natural environment. At first glance, the Mrt phenotype would seem to be the manifestation of context-dependent mutations, which are neutral in the wild, and only become deleterious in the lab environment. However, that scenario requires bidirectional mutation, such that Mrt alleles mutate into wild-type alleles as well as the reverse; if not, the population would eventually mutate its way to fixation for the Mrt phenotype. Bidirectional mutation is possible, of course, but the evidence at hand suggests it is not common, because the Mrt phenotype is associated with loss-of-function mutations.

A straightforward alternative to context-dependent neutrality is that Mrt alleles are deleterious in nature, in which case genetic variation is maintained by mutation-(purifying) selection balance (MSB). That possibility is intuitively attractive because, all else equal, sterility will never be favored by natural selection. All else may not be equal, however; the Mrt phenotype may be a pleiotropic correlate of some other trait(s) for which variation is maintained by some form of balancing selection.

To begin to sort out the various possibilities by which variation for the Mrt phenotype is maintained, we need to know (1) the rate of input of new genetic variation by mutation, and (2) the frequency of the Mrt phenotype in nature. At any one locus, the equilibrium frequency of a deleterious allele at MSB, 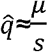, where μ is mutation rate from wild-type to the deleterious allele and *s* is the strength of selection against the mutant allele (the homozygous effect in an organism with near-complete self-fertilization, such as *C. elegans*; (Haldane 1927)). In a (nearly) completely inbred population, by extrapolation over the entire genome, the probability that an individual is not 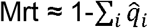, summed over all *i* loci capable of yielding the Mrt phenotype when mutated. In a population at MSB, the expected frequency of the Mrt phenotype is approximately 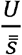, where *U* is the genome-wide rate of mutation to Mrt alleles and 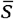 is the average strength of selection against an Mrt allele.

We estimated the rate of mutation to the Mrt phenotype from two sets of *C. elegans* laboratory mutation accumulation (MA) lines, which evolved in the near-absence of natural selection for approximately 250 generations. On average, each MA line carries about 65 unique spontaneous base-substitution and small indel mutations (Rajaei et al. 2021), and probably a few larger structural variants (A. S. Saxena and Baer, unpublished results). In addition, we estimated the frequency of the Mrt phenotype in a worldwide collection of 95 wild isolates. From these data, we infer the approximate strength of purifying selection acting on new Mrt mutations. Although we refer to “the mortal germline” as if it was a discrete, presence/absence trait, in reality, the mortal germline exists along a continuum (Frezal et al. 2018), and our analysis takes that fact into account.

## Materials and Methods

### Mutation Accumulation Experiment

Details of the mutation accumulation protocol are given in Baer et al. (2005). N2 is the standard laboratory strain of *C. elegans*; PB306 is a wild isolate generously provided by Scott Baird. The basic protocol follows that of Vassilieva and Lynch (1999) and is outlined in **Supplemental Figure S1**. Briefly, 100 replicate populations (MA lines) were initiated from a cryopreserved stock of a highly-inbred ancestor (“G0”) at mutation-drift equilibrium and propagated by transferring a single immature hermaphrodite at one-generation (four-day) intervals. Lines were maintained on 60mm NGM agar plates, spotted with 100 μ*l* of an overnight culture of the OP50 strain of *E. coli* B, at a constant 20°C. The lines were propagated for 250 transfers (Gmax=250), beginning in March, 2001 and culminating with a final cryopreservation in 2005.

### Wild Isolates

A collection of wild isolates of *C. elegans* was obtained from Erik Andersen (Northwestern University) in 2015 and cryopreserved in the Baer lab. A list of the wild isolates is given in **Supplemental Table S3**. The genome sequences of the wild isolates along with collection information are available at https://www.elegansvariation.org/.

### Mortal Germline Assay

The assay is based on that of Frezal et al. (2018), and is schematically depicted in **Figure 1**. Cryopreserved samples of the G0 ancestor were thawed onto 35 mm plates seeded with OP50 and incubated at 20° for three days, at which time 18 L4-stage hermaphrodites were picked individually to seeded 35 mm plates and incubated at 15°. The 18 replicates of the G0 ancestor were subsequently treated identically to the MA lines; we refer to these as “pseudolines” (PS). The following day, 35 randomly selected N2 (block 1) or PB306 (block 2) MA lines were thawed from cryopreserved samples and incubated at 15° C. PS and MA lines were allowed to reproduce for one generation (g.-3) at 15° C, at which point ten replicates of each line (MA and PS) were initiated by transferring a single L4 hermaphrodite to a seeded 35 mm plate. Each replicate was allowed to reproduce at 15°C for two more generations generation (g.-2, g.-1), at which time three L4s from each replicate were transferred to a seeded 35 mm plate (g.0) and incubated at 25° C. Subsequently, three L4s were transferred at three-day intervals for the duration of the assay. The N2 assay (block 1) was propagated for 21 generations; we terminated the PB306 assay (block 2) after 14 generations because it was evident that there were no lines with a strong Mrt phenotype.

**Figure 1.**
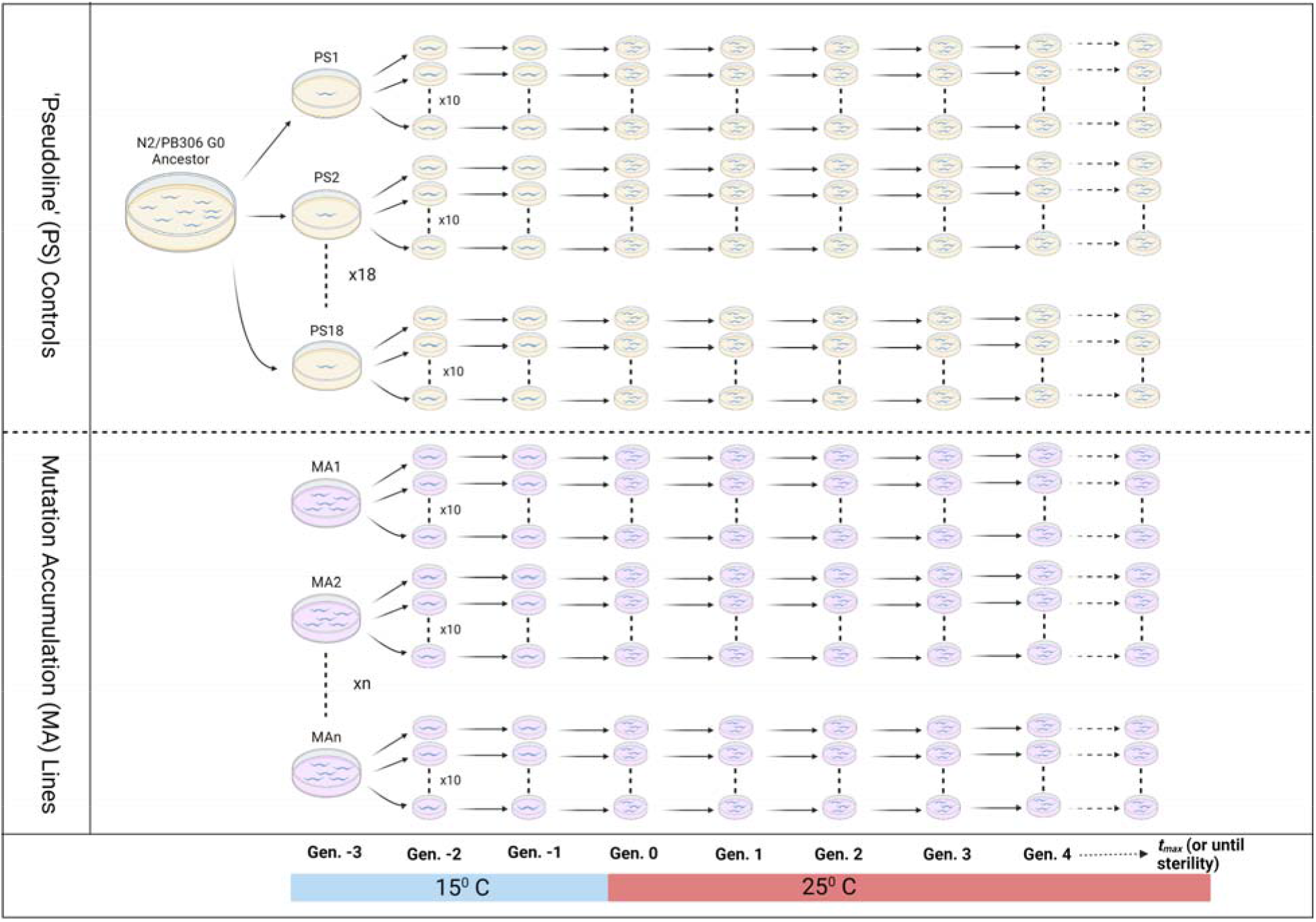
Schematic diagram of the Mrt assay. See Methods for details of the experiment.

The assay of the wild isolates was identical to that of the N2 MA lines (21 generations) except there were only three replicates per line rather than ten.

### The mortal germline (Mrt) phenotype

A strain is defined as having the Mrt phenotype if (1) it becomes sterile over a predefined number of generations (see next paragraph), and (2) sterility is manifested in a stereotypical way. Specifically, animals with the Mrt phenotype develop at approximately the normal rate, are of normal size and lifespan, exhibit typical activity, have a characteristically dark intestine, and an obvious absence of developing embryos (see Figure 1D of Frezal et al. 2018). Mrt sterility is defined in contrast to failure to reproduce *per se*. Some MA lines simply have low fitness, which may lead to failure to reproduce. Typically, worms from low-fitness lines are sickly-looking, develop slowly, mature at small size, and are sluggish. Individuals from low-fitness lines have low fecundity and/or lay eggs that fail to hatch, and often die before reproducing. Low fitness is not temperature-dependent, although the effects are often more severe at higher temperature (Matsuba et al. 2013). Several MA lines had low fitness; none of the 95 wild isolates did.

As noted, the Mrt phenotype exists along a continuum. We define a strain (MA or wild isolate) as having a “strong” Mrt phenotype if (1) the mean time to sterility is less than 10 generations, and (2) the maximum time to sterility is less than 15 generations. We further define a strain as having a “moderate” Mrt phenotype if (1) the mean time to sterility is less than 16 generations and (2) no replicate is still fertile by the culmination of the experiment at 21 generations. We define a “weak” Mrt phenotype as a strain that meets neither of the preceding criteria but in which at least two out of three replicates have become sterile by generation 21. A strain is designated as wild-type if at least two out of three replicates are fertile at generation 21. The strong-Mrt category is defined on the basis of the MA line results and to parallel the classification of Frezal et al. (2018); the moderate and weak categories are *ad hoc*.

### Haplotype tree

Mrt phenotypes as defined in the previous section were mapped onto a species-wide, whole-genome haplotype tree constructed from the *WI.20210121.hard-filter.isotype.min4* strain set, available from the CeNDR database (https://www.elegansvariation.org/data/release/latest). The tree was estimated by Neighbor Joining, as implemented in the QuickTree software (https://github.com/tseemann/quicktree).

#### Data Access

Mrt assay data are included in supplemental tables S1 and S3 and in Dryad ###. Genome sequence data of MA line 578 is deposited in the NCBI Sequence Read Archive under Accession number PRJNA665851, sample SAMN16272702.

## Results

### Mutation

Raw survival data are given in **Supplemental Table S1**. Both the N2 and PB306 progenitors are wild-type. In the N2 G0 progenitor, only ten of the 180 replicates (18 PS lines, 10 replicates/line) failed to reproduce before termination of the assay at generation 21, and only one of the 18 PS lines had more than one replicate fail to finish the assay. Of the 34 N2 MA lines assayed, one (line 540) incurred a heat-sensitive sterile mutation, identified as such because all ten replicates of the line were sterile after the first generation at 25°. Temperature-sensitive sterile and lethal mutations are well-documented in many organisms, and are distinct from Mrt. One line (line 578) had an obvious strong Mrt phenotype; all ten replicates were sterile by generation ten (median time to sterility = six generations). Of the remaining 32 MA lines, only two had more than one replicate fail prior to completion of the assay. Line 516 had 3/10 replicates fail, and line 538 had two. However, both lines had obviously low fitness (e.g. slow development, sickly worms) and did not exhibit the canonical Mrt-sterile phenotype, so we classify those lines as wild-type with respect to the Mrt phenotype.

Of the 180 replicates of the PB306 progenitor, only one failed to reproduce prior to completion of the assay at generation 14. Of the 33 PB306 MA lines, one (line 471) had 6/10 replicates fail before generation seven. However, line 471 has low fitness even at 20°, and the remaining four replicates survived to the end. The replicates that failed did not have the Mrt-sterile phenotype; rather, they were characterized by slow growth and dead worms. Accordingly, we do not classify line 471 as Mrt. No other PB306 MA line had more than one replicate fail to complete the assay.

From these data, we conclude that 1/34 N2 lines and 0/33 PB306 lines incurred a strong Mrt mutation, and no MA line incurred a moderate or weak Mrt mutation. We can calculate the point estimate of the genome-wide rate of mutation to (strong) Mrt as *U*_*Mrt*_=*k/nt*, where *k* is the number of Mrt mutations observed (one in N2 and zero in PB306, assuming that the one observed Mrt phenotype is the result of a single mutation, which it appears to be), *n* is the number of MA lines included, and *t* is the number of generations of MA. Note that this is the haploid rate, but that mutations accumulated in diploids; double the number of genomes (for diploidy) is cancelled by the probability of loss of a new neutral mutation in an MA line, which is 1/2. Pooling over the two sets of lines, the point estimate of *U*_*Mrt*_= 1/(67×250)≈ 6×10^−5^/genome/generation. If we assume that the number of mutations X is Poisson distributed among lines, the exact 95% confidence interval around *U*_*Mrt*_ can be calculated as follows. Let *λ*_*L*_ and *λ*_*U*_ be the lower and upper bounds on the (1-α)% confidence interval of a Poisson-distributed random variable X=*k*, defined as:

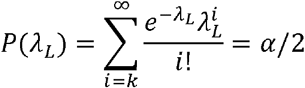

and

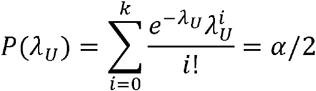

From the relationship between the Poisson and the Chi-square distributions, 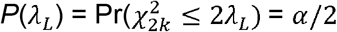 and 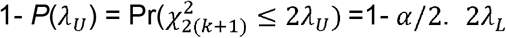 is the α/2 fractile of a *χ*^2^-distributed random variable with 2*k* degrees of freedom, and 2*λ*_*U*_ is the 1-*α*/2 fractile of a *χ*^2^-distributed random variable with 2(*k*+1) df (Ulm 1990). Here, *k*=1 mutation in *nt* (67 lines)(250 generations)=16750 meioses, so the 95% confidence interval around *U*_*Mrt*_ is (1.53×10^−6^ – 3.28×10^−4^/genome/generation). The per-nucleotide mutation rate in these lines is approximately 2.8×10^−9^/generation (Saxena et al. 2019; Rajaei et al. 2021) and the *C. elegans* genome is approximately 10^8^ bp, resulting in a point estimate of the mutational target of the Mrt phenotype of about 0.02%, and possibly as much as 0.1% of the *C. elegans* genome.

Given that one, and only one, MA line has a clear Mrt phenotype, we scrutinized its genome for candidate mutations (Rajaei et al. 2021; **Supplemental Methods**). Line 578 carries 37 unique base-substitutions, 13 deletions, and four insertions relative to the genome of the progenitor of the N2 MA lines (**Supplemental Table S2**). There is one obvious candidate, an 11-base frameshift insertion in an exon of the *nrde-2* gene. *nrde-2* is so-named for its <u>N</u>uclear <u>R</u>NAi <u>De</u>fective phenotype (Guang et al. 2008), is involved in heterochromatin assembly by small RNA as well as nuclear RNAi, and has been shown to be involved in temperature-dependent transgenerational nuclear silencing (Sakaguchi et al. 2014).

### Standing genetic variation

95 wild isolates (“strains”) were chosen haphazardly, based on a collection by E. C. Andersen. Assay data are given in **Supplemental Table S3**. Unlike that of the MA lines, the phenotypic distribution of the wild isolates cannot be unambiguously categorized into Mrt and Not Mrt. The difficulty has (at least) two sources. First, the sample size per strain is smaller (three replicates per strain, as opposed to ten per MA line), and second, there appear to be small-effect QTL segregating in the population that contribute a non-trivial fraction of the heritable variation (Frezal et al. 2018).

Six of the 95 strains had an unambiguous strong Mrt phenotype (**Figure 2**). Progressively loosening the Mrt criteria, 10 strains had the moderate Mrt phenotype, and another 13 strains had the weak Mrt phenotype. The remaining 65 strains were classified as wild-type, of which 42 remained fertile at 21 generations in all three replicates. The quantification is obviously not exact; some lines with relatively low mean time-to-failure were classified as wild-type because one replicate became sterile early on, whereas two of the three replicates remained fertile at 21 generations (e.g., EG4349). Depending on the stringency of the criteria, the frequency of the temperature-dependent Mrt phenotype in the wild isolates is at least 6/95 (~6%) and probably much higher.

**Figure 2.**
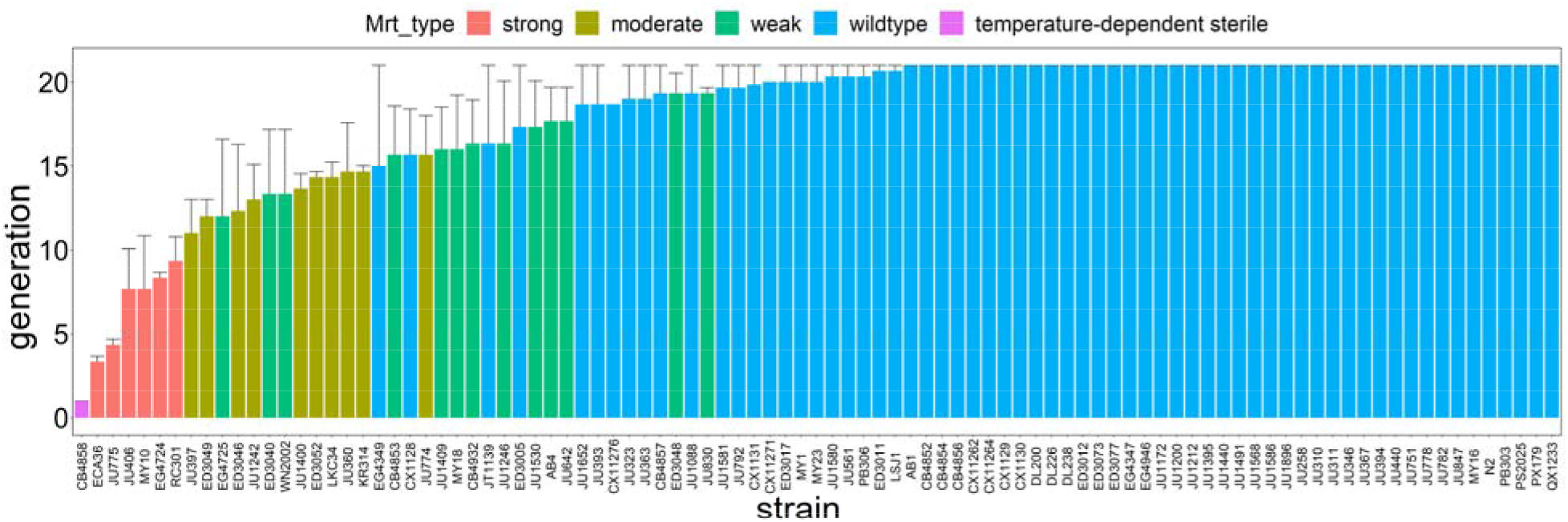
Average time to sterility of wild isolates (n=3 reps/isolate). Error bars are 1 SEM. See Methods for description of Mrt classification (Mrt_type).

### Mutation, selection, and the maintenance of genetic variation

We begin with the strong Mrt wild isolates, of which there are six. These strains clearly have the same strong Mrt phenotype as MA line 578. Solving the equation 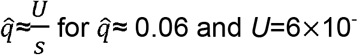 and *U*=6×10^−5^/generation, we infer a strength of selection *s*≈0.001. That strength of purifying selection is on the order of that inferred to be acting against deleterious alleles that affect competitive fitness (Yeh et al. 2017). The frequency of the strong Mrt phenotype is entirely consistent with genetic variation being maintained by MSB. The strength of selection against heat-sensitive sterile mutations can be similarly inferred, since one MA line (540) incurred such a mutation, and one of the 95 wild isolates (CB4858) had a heat-sensitive sterile phenotype, leading to an estimated selection coefficient *s*≈0.006.

The MA data are not so clear with respect to weaker Mrt phenotypes. The distribution of mean time-to-sterility among MA lines is no different from that of the G0 PS lines in either MA background (**Figure 3**). An upper 95% confidence limit on the genome-wide mutation rate to weak Mrt alleles that is consistent with observing no MA line with that phenotype can be calculated as before from the Poisson probability of observing *k* mutations, where now *k*=0. *U* as high as 2.2×10^−4^/generation is consistent with the observed absence of weak Mrt phenotypes among the MA lines.

**Figure 3.**
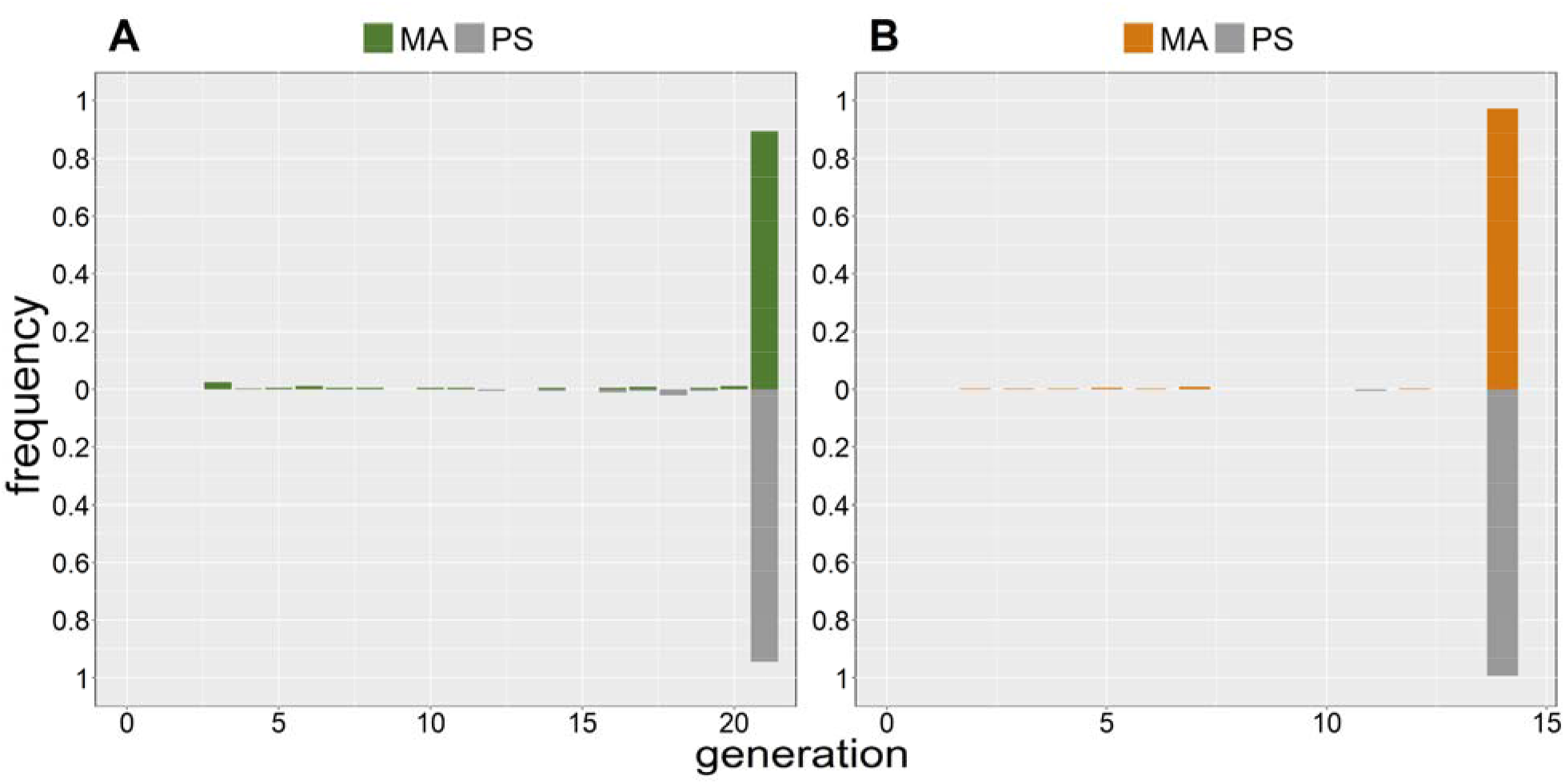
Frequency distribution of time to failure to reproduce of individual replicates in the MA assay. MA lines above the mid-line, G0 pseudolines (PS) below. (**A**) N2. (**B**) PB306.

A trait for which variation is maintained by MSB will recur and quickly be lost. When mapped onto a species’ haplotype phylogeny, the trait is expected to be scattered on tip branches throughout the tree, but not be present deeper in the tree. In contrast, a trait for which variation is maintained by balancing selection will appear deep(er) in the tree. We mapped the four categories of Mrt (strong, moderate, weak, and wild-type) onto a recent haplotype phylogeny of *C. elegans* (Supplemental Figure S2). The data are sparse, but for four of the six strong Mrt strains, the nearest neighbor in the tree with a characterized phenotype is wild-type, as predicted for a trait at MSB; the other two are ambiguous. The pattern is less clear for weaker Mrt phenotypes, but some clades do have multiple moderate and weak-Mrt strains, albeit interspersed with wild-type strains.

## Discussion

The high frequency of weak Mrt phenotypes in the wild isolates seems incongruous with the failure to observe even a single MA line with a weak Mrt phenotype. Long-term maintenance of neutral variation requires bidirectional mutation. It is evident that the strong Mrt phenotype is deleterious in the lab environment. Fortuitously, we have a strain (XZ1516) in long-term mass culture at 20° that has a strong temperature-dependent Mrt phenotype that is weakly penetrant at 20°. As an *ad hoc* test for back-mutation of a strong Mrt phenotype in the presence of selection, we initiated ten replicates of our Mrt assay at 25° with XZ1516 worms that had been cryopreserved after ~80 generations in mass culture at 20°. All ten replicates were sterile by seven generations. Obviously that would have been a more meaningful test had we kept the strain in mass culture at 25° rather than 20° (and used more than a single strain), but it at least suggests that back-mutations from a strong Mrt phenotype are infrequent. Of course, the genetic basis underlying weak Mrt phenotypes is likely to be different from that of the strong Mrt phenotype, which has been shown to typically result from loss-of-function mutations at protein-coding loci (e.g., MA line 578). Based on what is known about quantitative traits in general (Manolio et al. 2009; Boyle et al. 2017), it seems likely that much variation in weak Mrt is the result of variation in the magnitude and/or timing of expression of genes that confer a strong Mrt phenotype when silenced. On the other hand, "typical" quantitative traits accumulate abundant mutational variance (Houle et al. 1996; Davies et al. 2016), which is not the case for the weak Mrt phenotype in these lines. Another possibility is that a different epigenetically-heritable factor (e.g., a different small RNA) accumulates in the germline at a slower rate, leading to what we classify as a weak Mrt phenotype. The failure to observe a weak Mrt phenotype in the MA lines is consistent with that possibility.

Taken together, abundant genetic variation in nature coupled with a low rate of input of variation by mutation points toward variation being maintained by some type of balancing selection. We do not yet know enough about the natural history of the Mrt phenotype to identify candidate mechanisms, except to note the close correspondence between genes that produce a Mrt phenotype and the RNAi mechanism. Natural targets of RNAi include transposable elements and viruses (Robert et al. 2005; Fischer et al. 2013), each of which could plausibly constitute an agent of balancing selection.

## Acknowledgments

We thank José Miguel Ponciano, Matt Rockman, and the anonymous reviewers for their helpful comments. We especially thank Marie-Anne Félix for her comments and for piquing our interest in the mortal germline phenotype. Support was provided by NIH award GM127433 to CFB and V. Katju.

