## Supplemental Figure S2 for "Mutation, selection, and the prevalence of the *C. elegans* heat-sensitive mortal germline phenotype"

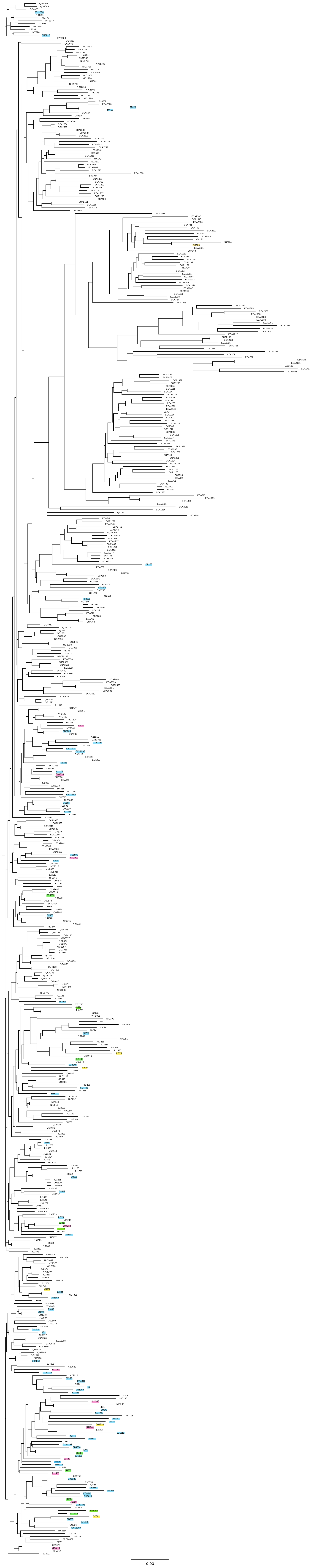
0.03

**Figure S2**. Neighbor-Joining tree of *C. elegans* whole-genome haplotypes. Strains highlighted in yellow are strong Mrt phenotype, green are moderate Mrt, purple are weak Mrt, and blue are wild-type. See Methods for details of phylogeny reconstruction and description of Mrt phenotypes.
