## Supplemental Methods for "Mutation, selection, and the prevalence of the *C. elegans* heat-sensitive mortal germline phenotype"

DNA Extraction and library preparation. Cryopreserved tubes of worms were thawed and grown for several days on 60 mm NGMA agar plates containing Streptomycin and Nystatin and seeded with 110 μl of overnight culture of the HB101 strain of *E. coli*. When food was nearly exhausted, a small piece (~50 mm^2^) of the agar plate was transferred onto a 100 mm NGMA plate and worms were allowed to grow for another 2-3 days. When food was nearly exhausted, worms were collected into 15 ml centrifuge tubes on ice and allowed to settle. Approximately 100 μl of settled worms were transferred into a 1.5 ml microcentrifuge tube and stored at -80°C until all samples were ready for DNA extraction, at which time samples were thawed and genomic DNA was extracted with the DNeasy Kit (Qiagen) following the manufacturer's protocol. Extracted DNA was stored at -20°C prior to library preparation.

Frozen DNA samples were thawed and diluted with diH_2_O to a concentration of 0.2 ng/μl. Tagmentation was performed per manufacturer’s instructions (Illumina, Nextera DNA Sample preparation kit, FC-121-1030). Following tagmentation, amplification was performed using custom barcoded IDT primers. Following amplification, a 96-pooled well sample was generated. Once pooled, 170µL of sample was combined with 30µL of 6× loading dye and run on a 2% agarose gel. DNA segments of 300-500bp were excised from gel. The QIAquick Gel Extraction Kit (QIAquick, 28704) was used to clean and elute DNA from the gel, following manufacturer’s instructions. Pooled library samples were quantified using the Qubit HS kit (Qubit, Q33230) and sequenced on a NovaSeq 6000 by Novogene, Inc.

Variant calling. Adapter sequence was trimmed from raw sequencing reads using fastp (Chen et al. 2018). Following trimming, we used bowtie2 (Langmead et al. 2009) to align trimmed sequence data to the N2 reference genome (WS263). Reads with the MQ<3 were removed from further analyses. Duplicate reads were identified and removed using MarkDuplicates tool in Genome Analysis Toolkit (GATK)/picard. Variants (SNPs and indels) were called using HaplotypeCaller (in BP_RESOLUTION mode) in GATK4 (v4.1.4.0) (McKenna et al. 2010). Next, the resulting files from all samples (23 *mev-1*, 68 N2, and 67 PB306 samples plus their ancestors) were consolidated using GenomicsDBImport, and variants called jointly using GenotypeGVCFs in GATK.

Variants that met the following three criteria were considered as putative mutations: (1) they were called homozygous; (2) they were present in one and only one MA line for each datastore, and (3) the ancestor genotype is homozygous wild-type (Saxena al., 2019). We initially applied a 3x coverage threshold for filtering the sites with <3x coverage and include only sites that are covered >3× in > 90% of the MA lines in each strain (for example 62 out of 68 in N2 MA lines). Putative indels ≥ 20 bp and all indels within 50 bp of another putative indel were visually inspected using the Integrative Genomics Viewer (IGV) software.

Potential functional effects of mutations were assessed using snpEFF 4.3 (Cingolani et al. 2012), and the annotation database WBcel235.82. Each variant potentially has multiple effects; snpEFF sorts them by potential impact (i.e., the highest putative impact is listed first). We include only the largest potential effect of each variant (**Supplemental Table S2**).
